# SteadyCellPhenotype: A web-based tool for the modeling of biological networks with ternary logic

**DOI:** 10.1101/2021.08.10.454961

**Authors:** Adam C. Knapp, Luis Sordo Vieira, Reinhard Laubenbacher, Julia Chifman

## Abstract

**Summary:** We introduce *SteadyCellPhenotype*, a browser based interface for the analysis of ternary biological networks. It includes tools for deterministically finding all steady states of a network, as well as the simulation and visualization of trajectories with publication quality graphics. Stochastic simulations allow us to approximate the size of the basin for attractors and deterministic simulations of trajectories nearby specified points allow us to explore the behavior of the system in that neighborhood.

**Availability:** https://github.com/knappa/steadycellphenotypeMITLicense

## 1 Introduction

Ternary bionetworks are generalizations of Boolean networks which allow variables to take three, rather than two, states. These have the advantage that they can model levels of a node being at a low, normal, or high state rather than simply being activated or not. In this sense, ternary networks lie in the middle ground between Boolean networks and ODEs, being more descriptive but keeping the qualitative nature of a Boolean network. Some example of ternary biomodels are Espinosa-Soto *et al.* (2004); *Remy et al*. (2015); Chifman *et al.* (2017).

The software landscape for Boolean networks has many useful tools for analysis and visualization (e.g. Naldi *et al.* (2009); Müssel *et al.* (2010); *von Kamp et al*. (2017); *Zheng et al*. (2009)). However, large ternary networks with arbitrary local activation functions cannot be easily analyzed by available software, which necessitated the creation of a reusable tool freely available to the biological community. *SteadyCellPhenotype* makes an important contribution to the landscape for ternary networks by adding several specialized features, that are to our knowledge not available in other packages. Our software can find *all* fixed points analytically without exhaustive search. For cyclic attractors, we take advantage of modern multi-core processors allowing researchers to quickly obtain stochastic samples. Other features include knockout/overexpression experiments and continuity, which is unique to non-Boolean networks. More importantly, our intuitive software allows the non-technical user to obtain results in a short time with multiple publication quality visualizations of the attractors and trajectories. On the other hand, advanced users have an option of command-line usage and the ability to add their own procedures.

*SteadyCellPhenotype* supports ternary networks in either polynomial form, through a collection of utility functions MAX, MIN, and NOT, and via combinations of both. The documentation for the allowed syntax, ternary networks in general, and all features is available at https://steadycellphenotype.github.io/.

## 2 Attractors and stochastic simulations

In biological contexts, attractors of discrete systems may correspond to distinct phenotypes Kauffman (1969). Our software uses synchronous updates, so each state belongs to the basin of a single attractor (fixed point or cyclic). Fixed points are of particular importance and our software is capable of computing all of them using Macaulay2 Grayson and Stillman (2021). We have tested large models of up to 64 nodes with running time approximately 2.6 min on a 2019 MacBook Pro (2.6Ghz, 6 core Intel i7). To find cycles, we convert the system into a NumPy Harris *et al.* (2020) function, which is dynamically compiled using the Numba library Lam *et al.* (2015), we then generate a user selectable number of random initial states and evolve them until they reach a limit cycle. For each path, we collect the number of iterations until we reach the limit cycle, see Figure 1a. This is of interest as a long winding path suggests that it may take some time for a cell to arrive at its phenotype.

**Fig. 1.**
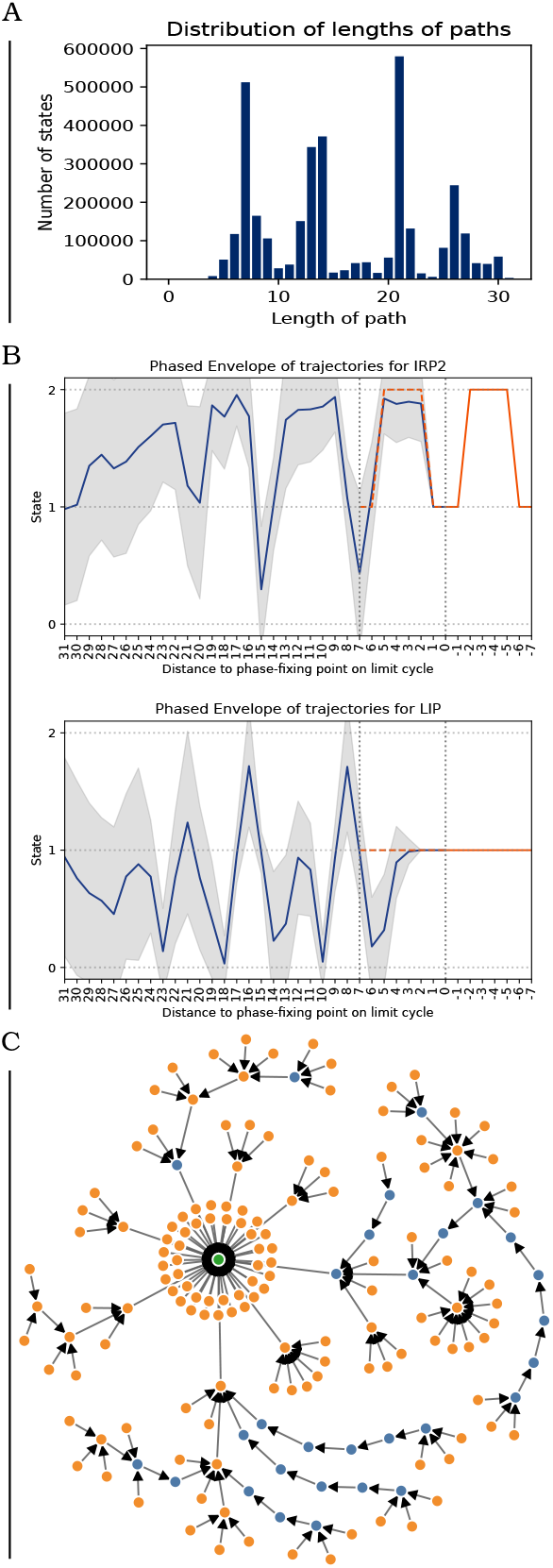
All figures produced using the model in Chifman et al. (2017) (A) Path lengths for *N* = 10^8^ random starts leading to fixed point. Spiky distribution indicates the presence of interesting topology in state transition graph. (B) Envelope for variables IRP2 and LIP indicate partial cyclic behavior beings before actual entrance to a 7-cycle. (C) States nearby the fixed point show disparate numbers of transitions to return to the attractor.

We also record statistics on the values of variables along the path. Paths are aligned “on the right” so that their arrival at a fixed point or a specific point on their cyclic attractor is simultaneous. The mean value and standard deviation of the variable states along these paths are shown in Figure 1b. This allow users to visualize if cyclic behavior was present in the system before the trajectory settled into the attractor or if the system exhibits transient organized behavior for multiple random initial conditions.

### Tracing a state and its neighbors

*SteadyCellPhenotype* also includes features for the analysis of trajectories from a single or collection of states. We provide a “trace states” feature to explore the evolution of a particular interesting state(s) and optionally the trajectories and attractors of nearby states, defined by having Hamming distance one from the selected state. The trajectory of these states are then computed and assembled into a directed graph, displayed using D3.js Bostock *et al.* (2011).

In Figure 1c we have a model with a single fixed point (in green) where we have assembled the trajectories of all of its nearby states. This graph tells us interesting information: while all nearby states return to the same fixed point, some of the trajectories are long. In biological contexts, this could mean that when one species is transiently perturbed from its original state, it takes the cell multiple time steps to return to this particular phenotype.

## 3 Conclusions

We believe *SteadyCellPhenotype* to be a useful tool for advancing the use and understanding of ternary networks. *SteadyCellPhenotype* is efficient, easy to use, and provides the researcher with meaningful data and publication quality visualizations not provided by other commonly used tools. Further work will include extensions to different numbers of states and a remotely hosted web platform.

